# Learned predictiveness acquired through experience prevails over the influence of conflicting verbal instructions in rapid selective attention

**DOI:** 10.1101/351908

**Authors:** Pedro L. Cobos, Miguel A. Vadillo, David Luque, Mike E. Le Pelley

## Abstract

Previous studies have provided evidence that selective attention tends to prioritize the processing of stimuli that are good predictors of upcoming events over nonpredictive stimuli. In the present study we explored whether the mechanism responsible for this effect critically reflects the influence of prior *experience* of predictiveness (history of attentional selection of predictive stimuli), or whether it reflects a more flexible process that can be adapted to new verbally acquired knowledge. Our experiment manipulated participants’ experience of the predictiveness of different stimuli over the course of trial-by-trial training; we then provided explicit verbal instructions regarding stimulus predictiveness that were designed to be either consistent or inconsistent with the previously established learned predictiveness. The effects of training and instruction on attention to stimuli were measured using a dot probe task. Results revealed a rapid attentional bias towards stimuli experienced as predictive (versus those experienced as nonpredictive), that was completely unaffected by verbal instructions. This was not due to participants’ failure to recall or use instructions appropriately, as revealed by analyses of their learning about stimuli, and their memory for instructions. Overall, these findings suggest that stimuli experienced as predictive through trial-by-trial training produce a relatively inflexible attentional bias based on prior selection history, which is not (always) easily altered through instructions.

## Introduction

Attention and predictive learning are intimately related in a bidirectional way. On the one hand, we learn more from attended stimuli than from unattended stimuli that are present concurrently in the environment [1–3]: That is, attention influences learning. On the other hand, learning about the *predictiveness* of stimuli has been shown to play an important role in determining how people subsequently allocate attention to those stimuli: That is, learning influences attention. A predictive stimulus is one that is a consistent and reliable indicator of the events that follow it, whether these events refer to presence of an outcome (e.g., electric shock) or its absence (no shock). A nonpredictive cue is one that provides no information regarding the events that follow it (e.g., a stimulus that is sometimes followed by shock, and sometimes by no shock). A wide range of studies has provided evidence consistent with the idea that people tend to allocate more attention to predictive stimuli than nonpredictive stimuli (see, for example, [1, 2, 4–6], for a review, see [7]).

Having established a relationship between learned predictiveness and attention, the next step is to determine the nature of the attentional process(es) that underlie this relationship.

One possibility is that the effect of predictiveness on attention reflects an effect of *selection history* [8, 9]. On this account, people learn that attending to predictive stimuli is advantageous, since it allows them to make accurate predictions about future events; in contrast, attending to nonpredictive stimuli is less useful since these stimuli do not allow accurate predictions to be made. As a result of this learning, people become more likely to select predictive stimuli than nonpredictive stimuli. Repeated experience of selecting predictive stimuli then induces an attentional bias towards these stimuli, which persists when they are encountered in future.

An alternative (though not mutually exclusive) possibility is that attentional biases towards predictive stimuli reflect the operation of relatively flexible attentional processes that are based on participants’ explicit knowledge regarding the current predictive value of stimuli, with this explicit knowledge arising through a process of inferential reasoning [10].

The critical difference between these two accounts relates to the information that drives the attentional bias. According to the selection history account, it is participants’ *experience* of the different utility of selecting predictive versus nonpredictive stimuli that determines the bias. On the ‘flexible processes’ account, it is participants’ explicit knowledge regarding the predictive status of stimuli that is critical; this explicit knowledge will be influenced by past experience of the consequences of selecting stimuli, but will also be influenced by verbally acquired knowledge independently of direct experience. This distinction thus raises the question: To what extent do effects of predictiveness on attention reflect an influence of experience (i.e., trial-by-trial training) versus an influence of verbal information? Below we review existing evidence on this issue that has produced mixed results, before describing a new experiment that aims to shed light on previous discrepancies.

The findings of a study by Mitchell et al. in 2012 [10] show that the allocation of attention to stimuli can be flexibly altered through verbal instructions. In their Experiment 2, participants underwent a first learning phase which established certain stimuli as predictive of the particular outcome that would occur on a trial, while other stimuli did not predict which outcome would occur (i.e., these latter stimuli were nonpredictive). Participants then completed a second learning phase during which all stimuli were paired with new outcomes. Importantly, immediately prior to this second phase, participants received instructions. Participants in the *Continuity* condition were told that those stimuli which had been predictive in Phase 1 would continue to be predictive in Phase 2, and those which had been nonpredictive would continue to be nonpredictive. Participants in the *Change* condition were told that stimuli which had been predictive in Phase 1 would now be nonpredictive, and vice versa. Mitchell et al. used eye-tracking to monitor overt attention to stimuli during Phase 2, in terms of dwell time – the length of time for which participants looked at each stimulus on each trial. Participants in the Continuity condition recorded longer dwell time on (i.e., attended more to) stimuli which had previously been predictive in Phase 1 than those which had been nonpredictive. In contrast, participants in the Change condition attended more to stimuli that had been non-predictive in Phase 1 than to stimuli that had been predictive. Judgments about the new stimulus-outcome relationships that were learned during Phase 2 also revealed that, in each condition, more was learned about the more-attended stimuli than about the less-attended stimuli.

Thus Mitchell et al.’s study [10] demonstrated a flexible influence of verbalisable knowledge on participants’ pattern of attention to predictive and nonpredictive stimuli. According to Mitchell et al.’s proposal, learners infer that stimuli that have been predictive of certain outcomes in a previous learning situation are also likely to be predictive of other outcomes in similar learning situations in future. This causal inference then leads learners to pay more attention to such stimuli through cognitive control processes that can be flexibly adapted on the basis of verbal instructions about the current predictive value of stimuli, in the absence of further training (i.e., trial-by-trial experience of predictive relationships). Mitchell et al. argued that their data suggested that attentional processes based on selection history, such as those envisaged by associative learning theories (e.g., [11, 12]), were unlikely to play any role in the effect of learned predictiveness on selective attention. This is because such theories would predict the prioritization, during Phase 2, of stimuli that had previously been *experienced* as predictive in Phase 1, regardless of verbal instructions (but see [13, 14], for conflicting evidence – an issue which we take up again in the General Discussion).

However, absence of evidence is not evidence of absence. It is possible that learning about predictiveness engages attentional processes based on both explicit knowledge and selection history, but that the particular measure of attention used by Mitchell et al. (gaze dwell time) was insensitive to the influence of selection history. Previous evidence suggests that effects of selection history are often relatively rapid and inflexible [7, 8]. Hence it remains possible that initial, rapid attentional orienting is influenced primarily by experience of predictiveness (i.e., by selection history), but that this initial experience-driven bias is subsequently overridden by a more flexible attentional control process based on explicit knowledge and reasoning – and it is this latter process that dominates in Mitchell et al.’s dwell time measure. Consistent with this possibility, Mitchell et al.’s dwell time measure summed gaze over a relatively long period (around 1 sec) and hence would be open to influence by relatively slow attentional processes. Moreover, whereas attention and eye movements are generally quite tightly coupled [15], it is possible for rapid shifts of attention to occur *covertly*; that is, in the absence of eye movements. Such covert attentional shifts would not be captured by eye-tracking. Thus it is possible that, even in the Change condition of Mitchell et al.’s study, there may have been a rapid (and possibly covert) attentional bias towards the stimulus that had been predictive during Phase 1, driven by selective history. This rapid bias may then have been followed by a second stage of overt attention to the previously-nonpredictive stimulus in line with verbal instructions that this stimulus would now be predictive, and it is this latter process that would be most evident in the dwell time measure used in this study.

In support of this explanation, recent evidence suggests that experienced predictiveness can indeed produce rapid attentional bias. Le Pelley, Vadillo, and Luque in 2013 [16] (see also [17]) trained participants on a task in which a pair of stimuli (coloured shapes)—known as a stimulus compound—appeared on each trial, with one stimulus on the left side of the screen, and the other on the right. Participants had to learn to make one of two button-press responses. One of the stimuli presented on each trial predicted the correct response, while the other was nonpredictive, much as in the study by Mitchell et al. [10]. However, in this case attention to the stimuli was measured using a *dot probe task* [18], which is based on the idea that detection of a target will be faster if that target appears in an attended location than in an unattended location.

On each trial of the dot probe task in Le Pelley et al.’s study [16], participants were shown (briefly) one of the stimulus compounds that had been experienced during training. After a short stimulus-onset asynchrony (SOA) of 250 ms, a dot (the probe) could appear at the location of one of the two stimuli. Participants were required to respond to the appearance of the probe as quickly as possible. Importantly, across trials of the test phase the probe was equally likely to appear in the location of (that is, be cued by) the stimulus that had been predictive during the training phase as it was to be cued by the nonpredictive stimulus. Hence there was no advantage to be gained in directing attention to either location prior to probe presentation. Indeed, participants were explicitly informed that in order to respond to the probe as quickly as possible, their best strategy was to ignore the initially presented stimuli.

Despite this instruction, responses to the probe were significantly faster when it was cued by the predictive stimulus than when it was cued by the nonpredictive stimulus, suggesting that participants had rapidly oriented their attention to the location of the predictive stimulus prior to the appearance of the probe. Notably, Le Pelley et al. [16] demonstrated that providing more time for participants to process the stimuli—by increasing the SOA on dot probe trials to 1000 ms—significantly *weakened* the influence of predictiveness on dot probe responding. Consistent with the argument that we advanced earlier, these findings demonstrate that rapid attentional biases that can be detected at short SOAs might go undetected in tasks that measure the deployment of attention over longer periods of time, including on the timescale of the measure used by Mitchell et al. (~1 sec).

In general terms, our hypothesis is that rapid attentional bias towards previously predictive stimuli is primarily determined by selection history, and relatively immune to the effect of instructions. To test this hypothesis, we conducted an experiment similar to Mitchell et al.’s Experiment 2 [10] but using a dot probe task (with short SOA) to measure rapid—and potentially covert—attention to stimuli. During Phase 1, some stimuli were trained as predictive of the correct categorization responses while others were nonpredictive. During Phase 2, participants learned new categorization responses. Immediately before this second phase, participants received continuity or change instructions regarding which stimuli would be important in determining the correct response in the following phase. A dot probe task was combined with the learning task throughout the experiment, as in Le Pelley et al.’s Experiment 3 [16] (see also [5, 19, 20]). By analyzing response times to the dot probe during Phase 2, we could examine the impact of experienced predictivess provided through training (in Phase 1) versus instructions on attentional bias. Crucially, in the change condition, we predicted an attentional bias driven by experienced predictivess within the short SOA condition. In other words, despite the conflict between experienced predictiveness and instructions regarding which stimulus should be prioritised, the former factor would have a greater influence on attentional bias than the latter.

## Materials and Methods

The design of our study was conceptually similar to that of Mitchell et al. [10] in that it compared the influence of training versus instruction on predictiveness-related attentional biases. Our study departed from the procedure of Mitchell et al. by using a within-subjects manipulation of verbal instructions, in order to increase the sensitivity of the experiment (a similar approach was used in Don & Livesey’s, Experiment 3 [13], and in Shone et al.’s Experiment 2 [14]). Accordingly, after Phase 1, participants were informed that four specific stimuli would be the most relevant to learn about during Phase 2. Participants then relevance were consistent with the predictive or nonpredictive status of stimuli that had been experienced during Phase 1 training (see Table 1). In contrast, in the *inconsistent pair*, instructions regarding relevance were inconsistent with the status of stimuli established during Phase 1. Finally, we also included two pairs of *novel* stimuli in Phase 2 that had not appeared in Phase 1. One stimulus of each pair was instructed as being relevant in Phase 2, whereas the other was not. Since these stimuli had not undergone prior training, any attentional bias revealed in dot probe responding for these novel pairs can only reflect the influence of instructions (cf. [21]). Observing an attentional bias for novel wthat participants had read, understood, and followed the instructions regarding relevance prior to Phase 2.

**Table 1.**
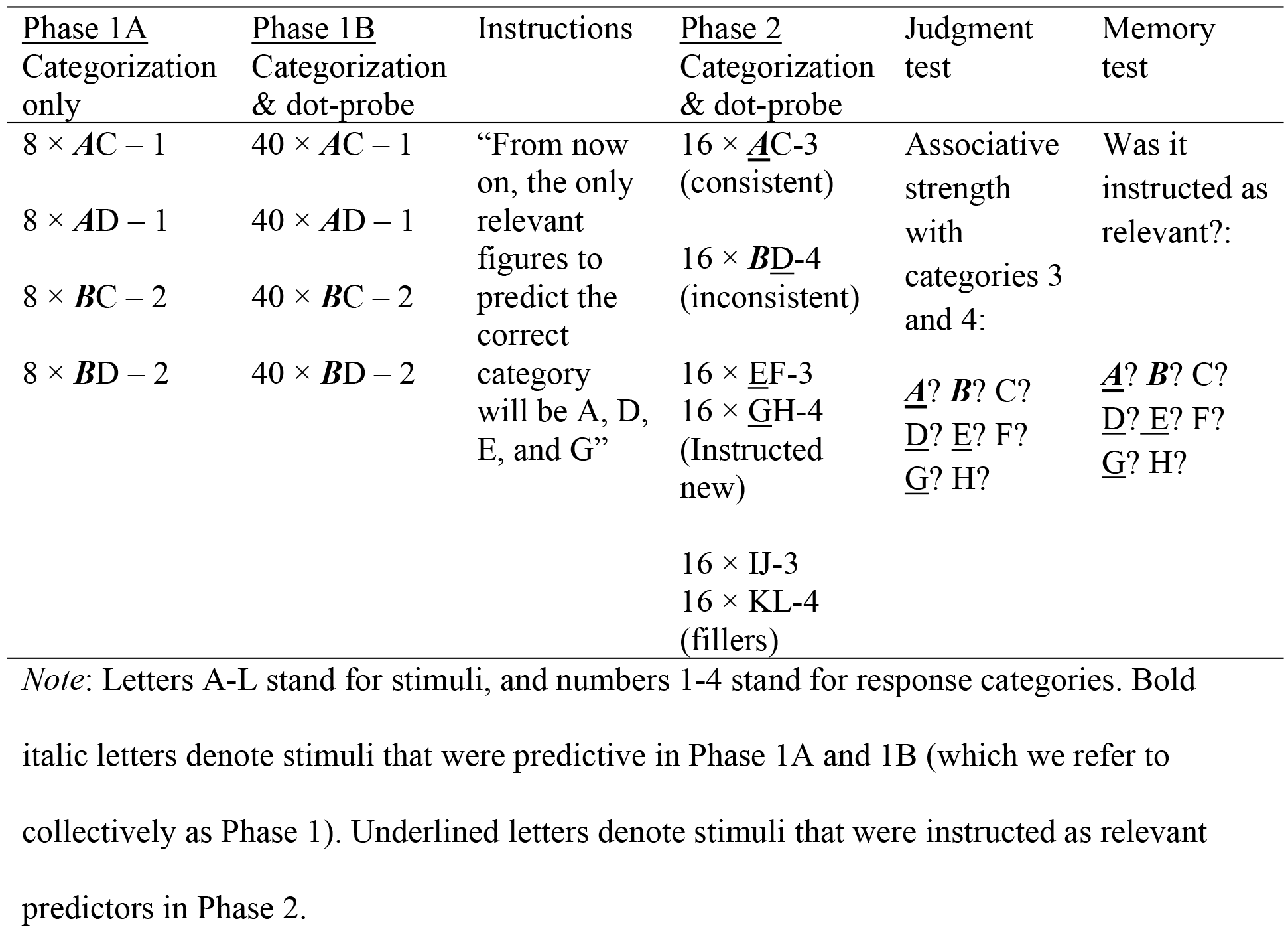
Design of the experiment.

### Participants and apparatus

A total of 135 students from a Spanish university participated for course credit; 68 were randomly assigned to a short SOA group, and the remainder to a long SOA group. Written consent was obtained and the Human Research Ethics Committee of the University of Malaga approved the study. The experiment was carried out in a quiet room with 10 semienclosed cubicles each equipped with a standard PC and 38.4 cm monitor. The task was run using the Cogent 2000 toolbox (http://www.vislab.ucl.ac.uk/Cogent/) for MATLAB. Participants made all responses with the computer keyboard.

### Stimuli

Stimuli were the same as those used by Luque et al. (2016), and included eight equal-sized circles (diameter subtending 4.7° visual angle at a viewing distance of ~80 cm), with radiating lines of varying thickness (see Fig 1). These figures were filled with different, easily discriminable colours that had similar brightness. The [red, green, blue] values for each colour were light red-brown [190, 86, 78], gold [190, 185, 78], green [93, 191, 77], turquoise [77, 191, 191], purple [132, 71, 255], pink [255, 5, 255], red [208, 0, 0], and grey [150, 150, 150]. These stimuli were randomly assigned the roles indicated by letters A-H in Table 1. Additionally, there were four more white outline figures consisting of two identical rectangles, one horizontally and the other vertically oriented, and two identical ellipses, one horizontally and the other vertically oriented. These last figures were used for filler trials, and were assigned roles corresponding to letters I-L in Table 1.

These stimuli were presented centrally in white square frames with sides subtending 6.4°, which were located on the right and left sides of a small fixation cross that was located in the centre of the screen; the centre-to-centre distance between the two boxes subtended 6.4°. The dot probe was a white square with side length subtending 1.1°. This would appear superimposed centrally on one of the stimuli. The screen background was black.

**Fig. 1.**
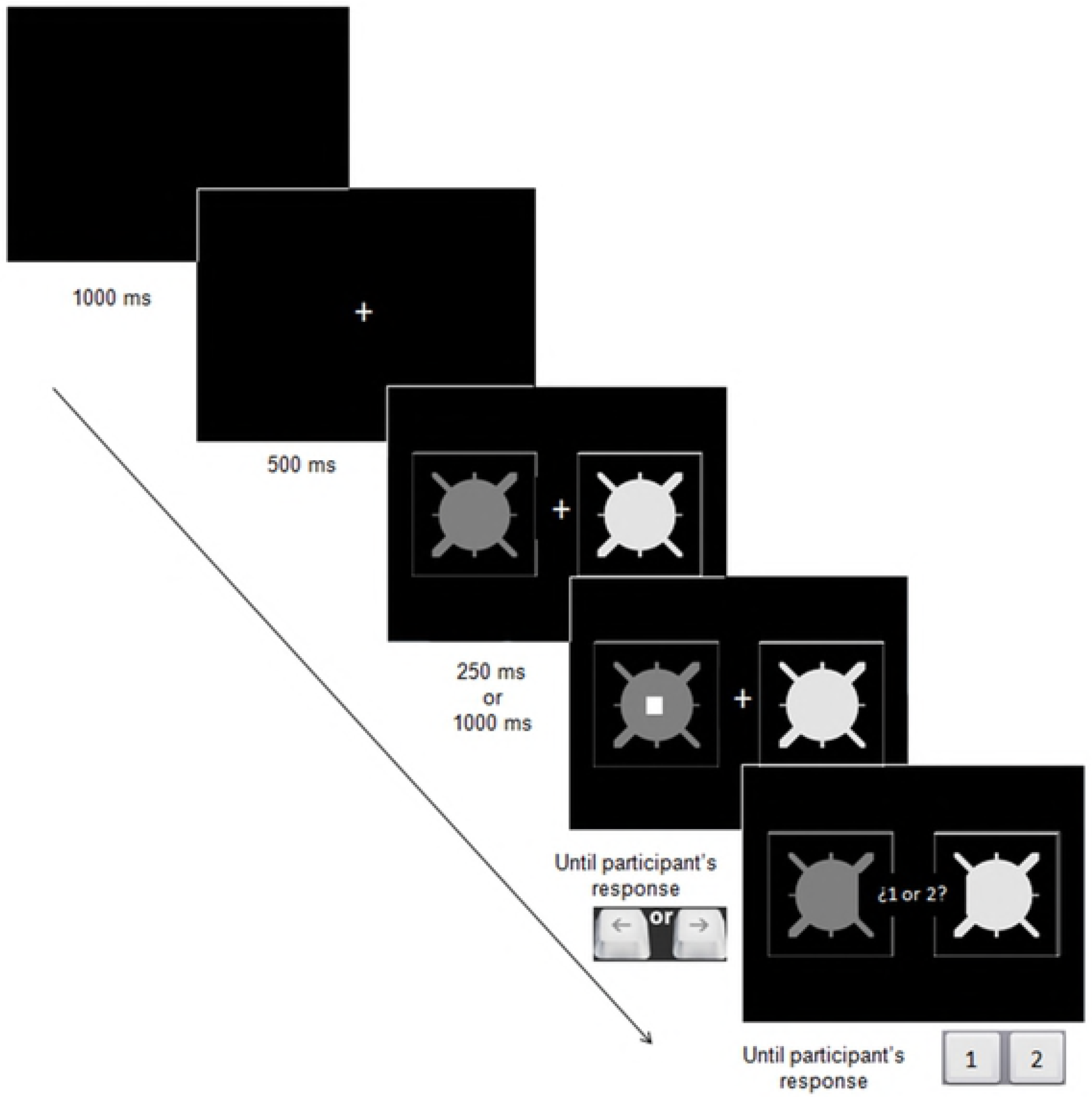
Stimulus display and timing of events on each training trial of the learning task.

### Procedure

The procedure was similar to that described in Le Pelley et al.’s Experiments 2 and 3 [16] (see also [5, 19]). Initial instructions (in Spanish) described the categorization task. Participants were told that, on each trial, a pair of stimuli would appear and they should make a categorization response by pressing either the ‘1’ or ‘2’ key with their left hand. Response keys ‘1’ and ‘2’ were randomly assigned the roles of response categories 1 and 2 shown in Table 1 for each participant. They were told they should try to learn the correct response for each pair of stimuli. Participants then underwent a first phase (Phase 1A) of 32 categorization trials. This comprised four eight-trial blocks, with each of the four stimulus pairs shown in Table 1 appearing twice per block in random order; for each stimulus pair, the predictive stimulus appeared once on the left and once on the right. On each trial a fixation cross appeared, followed after 500 ms by the pair of stimuli. After 1 s, a message framed within a central rectangle prompted participants to choose between response keys ‘1’ and ‘2’. Incorrect responses were followed by the feedback message “Error! The correct response was [1/2],” which remained onscreen for 3 s; no explicit feedback was provided for correct responses.

Following Phase 1A, participants received further instructions explaining that on subsequent trials they would complete two tasks: On each trial (a) a pair of stimuli would appear; (b) a small white square (the dot probe) would then appear superimposed on one of these stimuli; (c) participants should press the left or the right arrow with their right hand depending on whether the square appeared on the left or on the right stimulus, respectively; (d) once they had responded to the square, they should make a categorization response to the stimulus pair using the ‘1’ or ‘2’ keys with their left hand as in the pretraining stage. Participants were told that they should respond to the position of the dot probe as rapidly as possible and that “In order to do so, it is best that you ignore the pair of figures until you have responded to the location of the square” (translated from Spanish).

Fig 1 shows the event timing of a standard trial. Each such trial began with presentation of a central fixation cross. After 500 ms the stimulus pair appeared to either side of this cross. After an SOA of either 250 ms or 1,000 ms (depending on the SOA group to which the participant had been allocated), the dot probe appeared superimposed on one of the stimuli. This probe remained until participants made the correct response (left arrow key for a target presented on the left; right arrow key for a target on the right). Immediately on making the correct dot probe response, the probe disappeared and 1 s later the message “1 or 2?” appeared as for Phase 1A. Participants then made a categorization response using the ‘1’ or the ‘2’ keys; feedback was administered as in Phase 1A, and the next trial began after 1 s.

Participants completed Phase 1B, which comprised 10 blocks of 16 trials each (see Table 1). Each trial type of Phase 1B appeared four times; once for each combination of cue location (predictive cue on the left or on the right) and dot probe location (on the left or on the right stimulus). Therefore, the dot probe was equally likely to appear on the predictive or on the nonpredictive stimulus. The order of trials within each block was randomized.

Following Phase 1B, participants were told that in the next phase (Phase 2) they would learn new relationships between certain stimulus pairs and response categories 3 and 4 in a similar way as in Phase 1B. Some stimulus pairs had been presented in Phase 1A and 1B (which we refer to collectively as Phase 1), whereas others included new stimuli (see Table 1). Importantly, although all stimuli were in fact equally predictive of the response categories with which they were paired in Phase 2, participants were told that, from that moment on, the only relevant stimuli that they should use to choose the correct response key were A, D, E, and G. As explained in the Introduction, stimuli in Phase 2 were paired so as to create a consistent pair (AC) in which the instructed-relevant cue (A) had been predictive in Phase 1; an inconsistent pair (BD), in which the instructed-relevant cue (D) had been non-predictive in Phase 1; and two novel pairs (EF and GH), in which neither cue had appeared in Phase 1. Filler trials consisting of pairs IJ and KL were also included to increase the complexity of the learning task. The assumption underlying this procedural measure is that complex environments encourage the use of selective attention in order to focus and simplify information-processing. By increasing memory load in our critical test phase (Phase 2), we therefore hoped that these additional filler trials would provide additional drive for participants to deploy selective processes, e.g., by focusing on the cues mentioned in the verbal instructions. Phase 2 comprised four blocks of 24 trials each. Each of six stimulus pairs appeared four times per block, counterbalancing cue and probe location as in Phase 1B. Response categories 3 and 4 were randomly assigned to response keys ‘3’ and ‘4’ for each participant and independently of the assignment of response categories 1 and 2 to response keys ‘1’ and ‘2’. Thus, these assignments were uncorrelated across participants.

After Phase 2, participants completed a judgment phase in which they rated the extent to which each stimulus was associated with response categories 3 and 4, on a scale from 1 (‘completely sure that Stimulus X does not predict Response Y’) to 7 (‘completely sure that Stimulus X predicts Response Y’). Participants rated each stimulus with regard to each of the response categories (3 and 4) in random order.

Finally, participants completed a recognition memory test to assess their memory for the instructions regarding which stimuli were relevant and which were not. Again, a rating scale from 1 to 7 was used, with 1 meaning ‘completely sure that Stimulus X was not instructed as relevant, and 7 meaning ‘completely sure that Stimulus X was instructed as relevant’. Participants provided ratings for all stimuli in random order.

## Results

We imposed a selection criterion so as to exclude participants who did not show strong evidence of having learned the correct categorization responses. Specifically, we excluded data from participants who failed to reach a criterion of 80% correct categorization responses in the two last blocks of Phase 1B. This resulted in exclusion of five participants from the short SOA group (final *n* = 63), and eight from the long SOA group (final *n* = 59).

### Phase 1

Fig 2A shows the mean percentage of correct responses as a function of block and SOA group in Phase 1A (blocks 1-4) and 1B (blocks 5-14). Participants’ response accuracy increased over blocks; there was no apparent difference between SOA groups, with both approaching perfect accuracy during the final four blocks. These impressions were confirmed by a 14 (block) x 2 (SOA group: 250ms vs 1000ms) ANOVA, which yielded a significant main effect of block, *F*(13, 1560) = 99.5, *p* < .001, 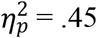. Neither the main effect of SOA nor the block x SOA interaction was significant (*F*s < 0.87). The same analysis within the last four blocks revealed a marginally significant effect of block, *F*(3, 360) = 2.16, *p* = .093, 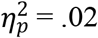. The main effect of group and the interaction between block and SOA were not significant (*F*s < 0.58). These statistical analyses yielded almost identical results even when the data from the excluded participants were included.

Following the same procedure as in Le Pelley et al. [16], response times (RTs) from the dot probe task were filtered and transformed before the analyses. First, RTs shorter than 150 ms and longer than 1500 ms were excluded, as were RTs from trials in which the first response to the probe was an incorrect response. Then, RTs were log-transformed to better fit a normal distribution. Transformed RTs lying more than 3 SDs from each participant’s mean were removed.

**Fig 2.**
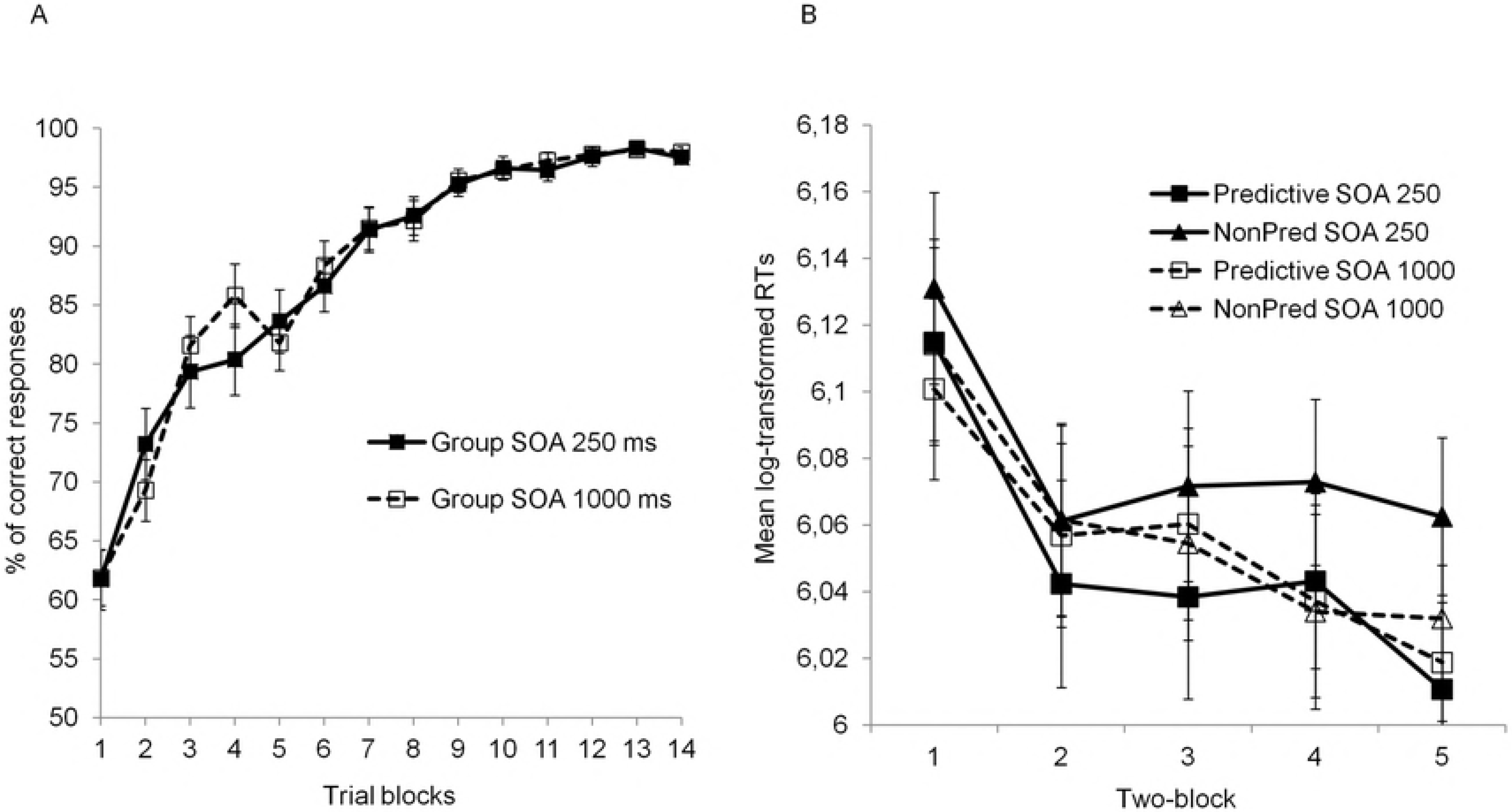
Summary of results found in Phase 1. Panel A: Mean percentage of correct categorization responses in Phase 1 as a function of SOA, group, and trial block. Panel B: Mean transformed response times to the dot in Phase 1 as a function of stimulus predictiveness, epoch, and SOA group (the dot probe task started in the fifth block of the learning phase). The intervals in both panels reflect the standard error of the mean.

Fig 2B shows mean log-transformed RTs as a function of dot probe position and SOA group, averaged over pairs of consecutive blocks (termed *epochs*). Participants in the short SOA group responded faster when the probe appeared on the predictive stimulus than when it appeared on the nonpredictive stimulus. This tendency was greater in late than in early epochs. In contrast, participants in the long SOA group showed similar RTs regardless of the probe’s position. A 2 (probe position: Predictive vs nonpredictive stimulus) × 5 (epoch) × 2 (SOA) ANOVA revealed main effects of probe position, *F*(1, 120) = 15.1, *p* < .001, 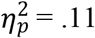, and epoch, *F*(4, 480) = 19.3, *p* < .001, 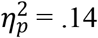, and a significant probe position × SOA interaction, *F*(1, 120) = 8.27, *p* = .005, 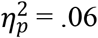 (*F*s < 1.5 for all remaining effects, smallest *p* = .202). A follow-up 2 (probe position) x 5 (epoch) ANOVA within the 250 ms SOA group yielded significant effects of probe position, *F*(1,62) = 19.56, *p* < .001, 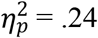, and epoch, *F*(4,248) = 8.98, *p* < .001, 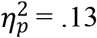, and a marginally significant interaction, *F*(4, 248) = 2.24, *p* = .066, 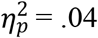. The same analysis within the 1000ms SOA group found only a significant effect of epoch, *F*(4, 232) = 11.94, *p* < .001, 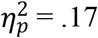; other *F*s < 1.

The results of the dot probe task essentially replicate Le Pelley et al.’s Experiment 3 [16], and indicate that predictive learning tended to produce an attentional bias towards the predictive stimulus. The fact that this bias was found in the 250 ms SOA condition but not in the 1000 ms SOA condition implicates a very rapid and short-lived attentional bias towards predictive stimuli.

### Phase 2

Fig 3A shows mean log-transformed RTs for ‘old’ stimuli A-D (i.e., stimuli previously experienced during Phase 1) as a function of experienced predictiveness, instructions, and SOA group, averaged across Phase 2. A 2 (experienced predictiveness: probe appeared on stimulus that had been predictive in Phase 1 vs stimulus that had been nonpredictive) × 2 (instructions: probe appeared on stimulus that had been instructed as relevant vs noninstructed) × 2 (SOA) ANOVA yielded a marginally significant effect of experienced predictiveness, *F*(1, 120) = 3.17, *p* = .077, 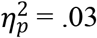 and a marginal experienced predictiveness × SOA interaction, *F*(1, 120) = 3.37*, p* = .069, 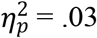 (other *F*s < 1.97, smallest *p* = .184). This interaction between experienced predictiveness and SOA is consistent with the results from Phase 1 and with Le Pelley et al. [16]. A follow-up 2 (experienced predictiveness) × 2 (instructions) ANOVA within the 250 ms SOA group yielded only a significant effect of experienced predictiveness, *F*(1, 62) = 6.61, *p* = .013, 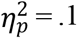 (other *F*s < 0.15). Similar analysis within the 1000 ms SOA group found no significant effects (*F*s < 0.5).

As expected, the short SOA group showed an attentional bias towards stimuli previously learned to be predictive through trial-by-trial training. Crucially, this effect was not significantly affected by whether these stimuli had been explicitly instructed as relevant or not during Phase 2. In contrast, the long SOA group showed similar RTs regardless of the experienced predictiveness of stimuli or instructions. The fact that an attentional bias towards predictive stimuli was detected at short SOA but not long SOA is again consistent with the engagement of a fast attentional process that may go undetected if the attentional task does
not impose strong enough time constraints.

**Fig 3.**
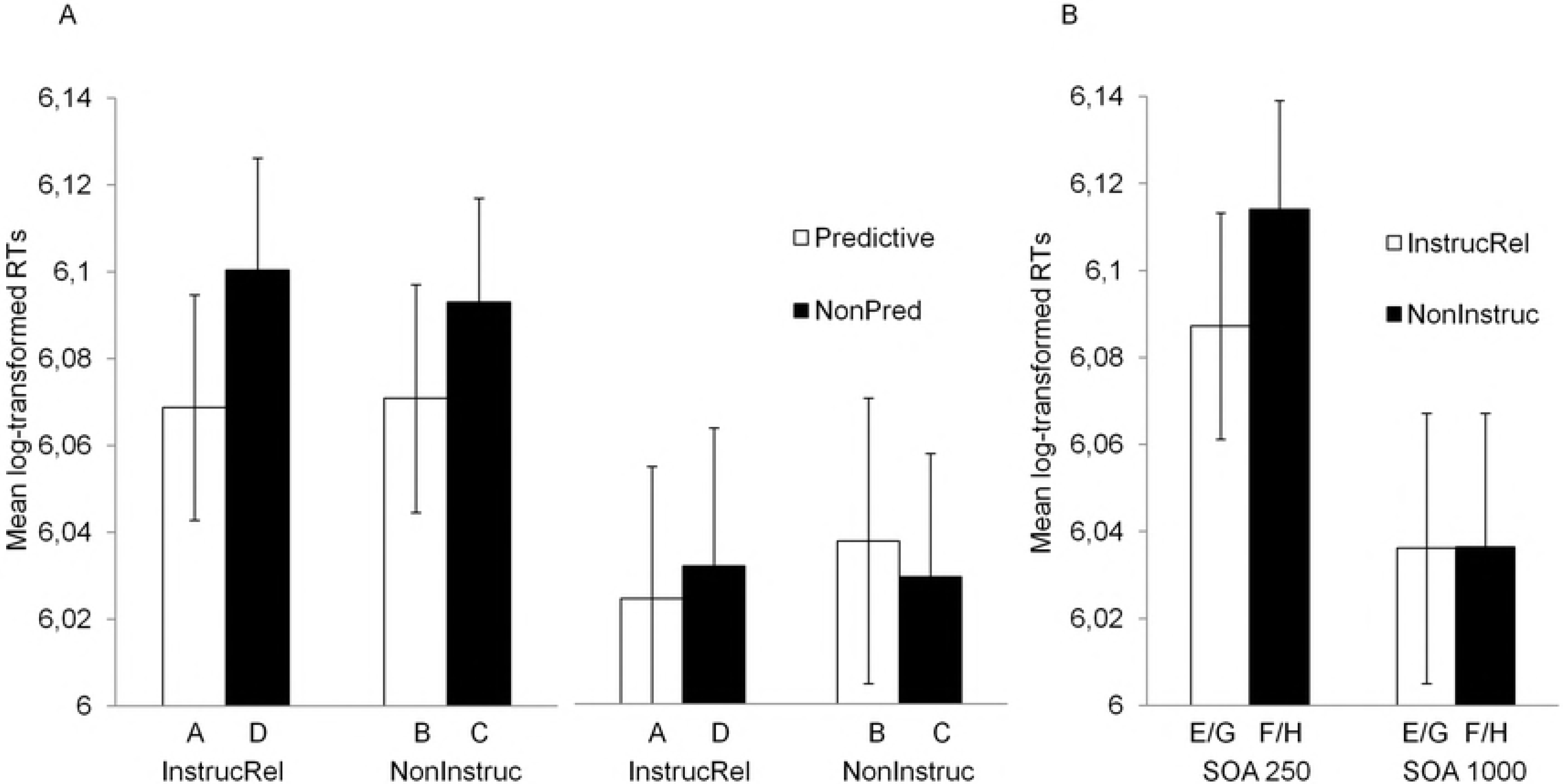
Results from Phase 2. Panel A: Mean log-transformed response times to the dot when it appeared on stimuli A-D, whose predictiveness had been established through previous experience in Phase 1. Results are displayed as a function of learned predictiveness, instructions regarding stimulus relevance, and SOA group. Panel B: Mean log-transformed response times to the dot when it appeared on new stimuli E-H, which did not form part of previous experience provided through Phase 1. Results are displayed as a function of instructions regarding stimulus relevance and SOA group. In both panels, intervals reflect the standard error of the mean.

One possible explanation of the failure of instructions to exert any significant effect on the data in Fig 3A is simply that participants did not read, understand, or make use of these instructions during Phase 2. To test this possibility, we analyzed the effects of instructions on RTs for novel stimulus pairs EF and GH. Recall that stimuli E and G were instructed as relevant during Phase 2, while F and H were noninstructed; none of these cues was experienced during Phase 1. Fig 3B shows mean log-transformed RTs during Phase 2. For these novel pairs, participants in the 250ms SOA group responded faster when the probe appeared on instructed stimuli than when it appeared on noninstructed stimuli. Participants in the 1000ms SOA group did not show a clear bias. A 2 (instructions: instructed vs noninstructed) × 2 (SOA), ANOVA yielded no significant effect (all *F*s < 2.76, smallest *p* = .1). However, since our previous analyses suggest that attentional biases in the dot probe task were confined to the short SOA group, we used a t-test to analyze the effect of instructions on RTs within the short SOA group only. This revealed a significant effect of instructions, *t*(62) = 2.33, *p* = .023, *η*^2^ = .08. This confirms that participants *were* effectively following the instructions about stimulus relevance, and that such instructions can produce a rapid attentional bias towards stimuli, at least when such instructions do not conflict with stimulus predictiveness experienced through trial-by-trial training.

### Ratings of stimulus-outcome relationships

Participants’ ratings from the Judgment Test were analyzed to assess the influence of experienced predictiveness and instructions on learning of stimulus-outcome relationships in Phase 2. Following Le Pelley and McLaren [3] (see also [22]), we calculated a rating score for each stimulus by subtracting the rating given to the incorrect response category from the rating given to the correct response category. High, positive values on this scale (maximum = 7) indicate strong learning of a correct stimulus-outcome relationship. Table 2 shows mean ratings for each stimulus (ratings for cues E and G, which were equivalent, were combined [denoted E/G]; ditto for cues F and H). The data relating to cues A-D were analyzed with a 2 (experienced predictiveness: Predictive vs nonpredictive during Phase 1) x 2 (instruction) x 2 (SOA) ANOVA. This revealed significant main effects of experienced predictiveness, *F*(1, 120) = 28.3, *p* < .001, 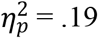, and instruction, *F*(1, 120) = 6.03*, p* = .015, 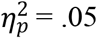. No other effects were significant (*F*s < 1.51, smallest *p* = .221). Both short and long SOA groups learned more during Phase 2 about stimuli that had previously been experienced as predictive than those that had been experienced as nonpredictive. Both groups also learned more about stimuli that had been explicitly instructed as relevant during Phase 2 than those that had not been instructed. This latter finding once again confirms that participants read and made use of the instructions regarding relevance given prior to Phase 2.

**Table 2.** Mean rating scores and standard deviations of the means (in parentheses) for stimulus-outcome relationships learned in Phase 2.

| SOA<br>group | Stimulus type |  |  |  |  |  |
| --- | --- | --- | --- | --- | --- | --- |
|  | Instructed relevant |  |  | Noninstructed |  |  |
|  | Predictive<br>(A) | NonPred<br>(D) | New<br>(E/G) | Predictive<br>(B) | NonPred<br>(C) | New<br>(F/H) |
| 250 ms | 4.37<br>(0.42) | 2.37<br>(0.5) | 4.25<br>(0.3) | 3.71<br>(0.45) | 2.19<br>(0.51) | 4.12<br>(0.3) |
| 1000 ms | 4.03<br>(0.44) | 2.59<br>(0.48) | 4.37<br>(0.29) | 3.42<br>(0.5) | 0.85<br>(0.57) | 3.86<br>(0.34) |

Putting together the dot probe results from Phase 2 and participants’ ratings for old stimuli, it seems that past experience with stimuli had an influence on rapid and short-lived attentional bias towards predictive stimuli, and on how much is learned about such stimuli in a subsequent phase of learning. Additionally, instructions about stimulus relevance had an effect on learning, as measured by subjective ratings, but not on rapid and short-lived attentional capture.

Regarding Stimuli E-H, there was a numerical trend towards higher ratings for the instructed cues than the noninstructed cues, but it did not reach statistical significance. A 2 (instruction) × 2 (SOA group) ANOVA on participants’ ratings revealed no significant effects (all *F*s < 1.55, smallest *p* = .217). Thus, in this case, instructions exerted an effect on attentional capture that did not translate into an advantage in terms of stimulus-outcome learning (as measured by explicit judgements).

### Recognition ratings

Fig 4 shows participants’ mean recognition ratings as a function of instructions, type of stimulus, and SOA group. The different groups of stimuli in the figure correspond to the compounds presented during Phase 2. This highlights the extent to which recognition memory was affected by the congruency between participants’ experienced predictiveness, and the instructions they received. A 2 (instructions) × 3 (stimulus type: AC vs BD vs EF/GH) × 2 (SOA) ANOVA on participants’ recognition ratings revealed significant main effects of instruction, *F*(1, 120) = 64.85*, p* < .001, 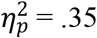, and stimulus type, *F*(2, 240) = 11.34, *p* < .001, 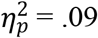, and an instruction × stimulus type interaction, *F*(2, 240) = 8.25, *p* < .001, 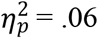 (other *F*s < 2.32, smallest *p* = .101).

**Fig 4.**
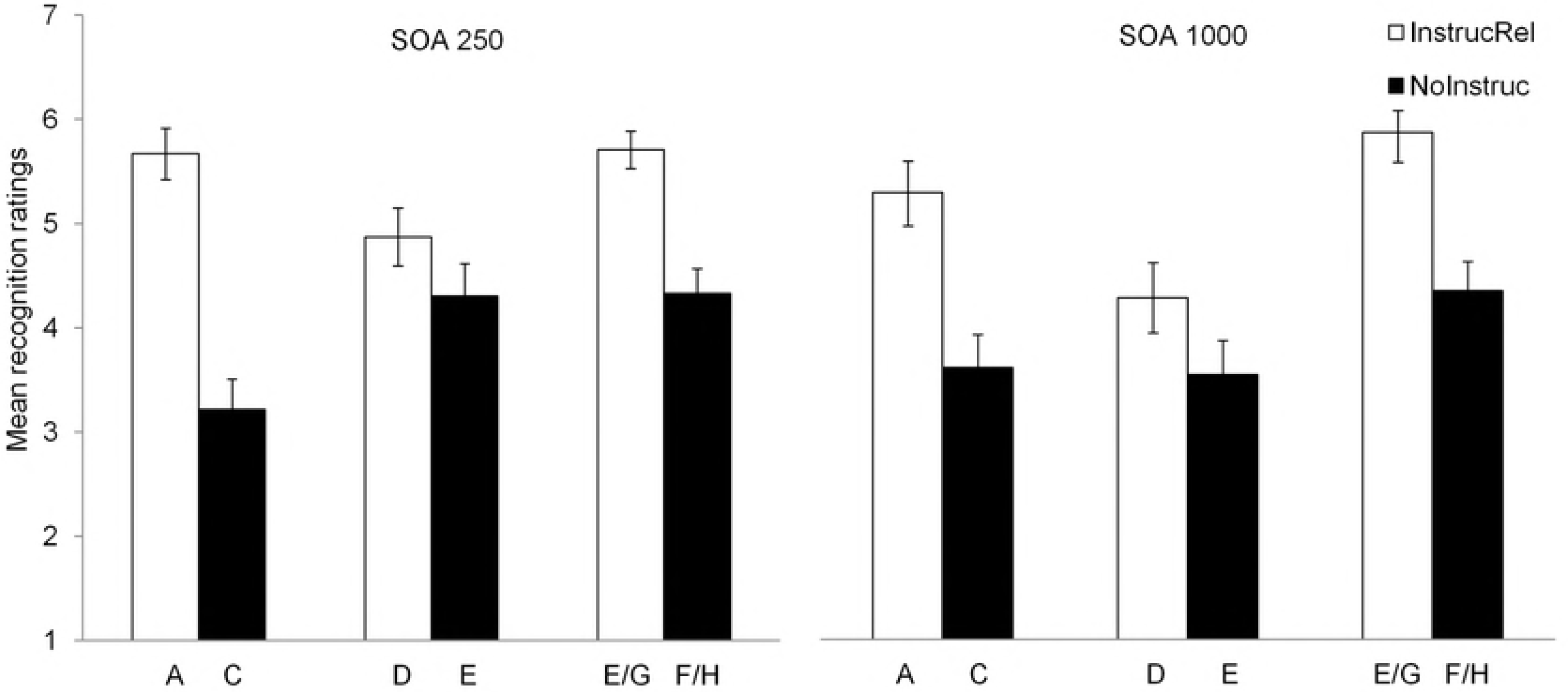
Mean recognition ratings. Mean recognition ratings for stimuli as a function of instructions regarding relevance, and SOA group.

Overall, recognition ratings were higher for cues that were instructed as relevant than for those that were not instructed, confirming again that participants had read and remembered these instructions. Interestingly, however, the effect of instructions differed as a function of stimulus type: Fig 4 suggests a larger effect of instructions for stimuli belonging to the *consistent pair* (AC) than for the *inconsistent pair* (BD), with an intermediate effect for stimuli belonging to novel pairs (EF and GH). Nevertheless, analysis of simple effects (collapsing across SOA groups) revealed a significant effect of instructions for each type of compound: for AC, *F*(1, 121) = 64.04, *p* < .001, *η*^2^ = .35; for BD: *F*(1, 121) = 4.69, *p* = .037, *η*^2^ = .04; and for EF/GH: *F*(1, 121) = 42.2, *p* < .001, *η*^2^ = .26.

## Discussion

We examined the influence of both prior training experience (selection history) and verbal instructions on predictiveness-driven attentional biases. To this end, participants experienced differences in the predictiveness of different stimuli over the course of trial-by-trial training in a first learning phase, and, later on, received verbal instructions regarding stimulus relevance for the subsequent learning phase that could be either consistent (AC compound) or inconsistent (BD compound) with experienced stimulus predictiveness. We measured the effects of these manipulations on spatial cueing in the dot probe task following the same procedure as Le Pelley et al.’ Experiment 3 [16]. Like Le Pelley et al. [16] (see also [17]), the current experiment found that—with a short stimulus-onset asynchrony (SOA) between the stimuli and the probe—responses to the probe during Phase 2 were faster when its position was cued by stimuli previously experienced as predictive compared with nonpredictive stimuli. This suggests that experienced predictiveness produced an attentional bias towards predictive stimuli. The fact that experienced predictiveness produced a bias in spatial cueing of the probe only at short SOA (250 ms) and not at longer SOA (1000 ms), suggests the operation of a rapid and short-lived attentional process.

Most importantly, the rapid attentional bias towards predictive stimuli (observed at short SOA) was not reversed or even significantly altered by conflicting verbal instructions regarding stimulus relevance. This was not due to participants’ failure to understand, retrieve, and follow verbal instructions. First, instructions regarding stimulus relevance affected explicit ratings about stimulus-outcome relationships learned in Phase 2. These ratings clearly show that participants tended to learn more about stimuli instructed as relevant (A & D) than noninstructed stimuli (B & C). Second, instructions produced an attentional bias towards new stimuli instructed as relevant (E & G) relative to new stimuli that were noninstructed (F & H). Finally, memory for instructions was reasonably good as evidenced by participants’ higher recognition ratings to stimuli instructed as relevant than noninstructed stimuli.

Thus, despite evidence that participants had read, understood, and implemented verbal instructions regarding stimulus relevance, these instructions had no effect on the bias in rapid attentional orienting to stimuli that had previously been *experienced* as predictive, compared to those experienced as nonpredictive. This suggests that trial-by-trial experienced predictiveness (i.e., selection history) drives the development of a rapid and relatively inflexible attentional bias that is somewhat insulated from changes in explicit knowledge about predictive status produced by verbal instructions. Note that we are not claiming here that performance in the dot probe task at short SOA is *generally* immune to verbal instructions. Indeed, our own data suggest this is not the case – for the novel stimuli (that had not been experienced during Phase 1), responses to the dot probe were significantly faster when it was cued by a stimulus that had been instructed as relevant (E/G) than when it was cued by a stimulus that had not been instructed (F/H) (for related findings, see [23, 24]). The novel finding of our data is that the influence of prior experience of predictiveness on rapid attentional bias is sufficiently strong that, given a difference in selection history, no effect of attentional control via instruction is observed.

In line with previous evidence [10, 13, 14], we found that participants’ learning of stimulus-outcome relationships during Phase 2 was influenced by instructions regarding relevance: Participants learned more, in general, about stimuli instructed as relevant than those that were not instructed. That said, the influence of instructions on learning was relatively slight, and was not sufficient to overcome the influence of experienced predictiveness on learning. That is, we also observed a main effect of experienced predictiveness on participants’ judgments of stimulus-outcome relationships, and instructions were not sufficient to reverse the pattern of greater learning about stimuli experienced as predictive than those experienced as nonpredictive. This is indicated by the finding that stimulus D (experienced as nonpredictive but instructed as relevant) produced weaker judgments than stimulus B (experienced as predictive but not instructed as relevant). Thus while demonstrating an influence of verbal instructions about stimulus relevance on learning, our data fail to replicate Mitchell et al.’s finding of a complete reversal of the effect of experience as a result of instructions [10]. In this respect our data are more similar to subsequent findings that have also failed to replicate this full reversal [13, 14]. Taken together, these findings suggest that both selection history produced via repeated experience with stimuli, and verbalisable knowledge, may contribute to biases in learning towards predictive cues observed in earlier studies (e.g., [2, 3, 25]).

It is noteworthy that a significant influence of instructions on stimulus-outcome judgements was observed only for stimuli that had previously been experienced during Phase 1 – no significant effect of instructions was seen for novel cues E-H. This pattern was unexpected: One might naturally expect that, in the absence of any other reason to attend to one stimulus or the other, participants would tend towards the stimulus instructed as relevant. It is unclear what to make of this null finding, and we note that there was a numerical trend towards greater learning about the instructed stimulus. One possibility is that the nonsignificant effect may reflect formation of a strong within-compound association between the elements of ‘new’ compounds EF and GH. For example, F was only ever experienced in compound with E, and hence a relatively strong association may have formed between these stimuli (compared to stimulus A for example, which was sometimes experienced with C and sometimes with D). Suppose that participants followed instructions regarding the relevance of new stimuli to the *outcome*, and learned a stronger stimulus-outcome association for stimulus E than stimulus F. Participants may still show strong responding to stimulus F on test, if presentation of F retrieves the memory of E (via the strong within-compound association), which in turn retrieves the outcome via the strong E-outcome association. In the absence of further evidence, however, this account is currently speculation.

We implemented instructions regarding stimulus relevance by explicitly informing participants which specific stimuli would be relevant during Phase 2 (following a procedure used by Don & Livesey in 2015 [13], and by Shone et al. in 2015 [14]). This differed from the approach used by Mitchell et al. [10], who provided the more general instruction that stimuli which had been predictive during Phase 1 were highly likely (in the Continuity condition) or highly unlikely (in the Change condition) to be predictive during Phase 2. It seems unlikely that this procedural difference was responsible for the persistent, rapid attentional bias towards stimuli experienced as predictive observed in the dot probe task of the current experiment. As Don and Livesey [13] noted, the instructions used by Mitchell et al. [10] might actually result in a rapid attentional bias towards stimuli previously experienced as predictive even in the Change condition, since participants may first need to identify the stimulus that was previously predictive in order to identify the stimulus which was previously nonpredictive (and which should now be attended, according to instructions). In contrast, direct instruction regarding which cues are relevant in Phase 2 does not require that participants first identify the stimulus which used to be predictive in Phase 1. Consistent with this claim, Don and Livesey [13] showed that instructing the relevance of specific stimuli results in, if anything, a *larger* influence of instructions on stimulus-outcome learning than does providing more general instructions regarding continuity/change, as used by Mitchell et al. [10]. This implies that the procedure used in the current experiment should have been at least as sensitive to showing an effect of instructions on attentional orienting as that used by Mitchell et al., if such an effect were to exist.

The primary aim of the current experiment was to assess whether—and the extent to which—the influence of experienced predictiveness on attention reflects the operation of processes based on selection history (modulated by experience) versus explicit knowledge (modulated by experience and verbal instructions). Our data suggest that both play a distinct role. In this final section, we briefly consider the nature of these attentional processes. One interpretation is that the rapid and short-lived influence of selection history reflects a relatively automatic process over which participants have little strategic control (cf. [8, 9, 26]). On this account, repeated experience of attentional selection of a particular stimulus produces an automatic and habitual prioritization of that stimulus. In the current dot probe task, the locations of the predictive/nonpredictive stimuli were noninformative with regard to the location in which the probe would appear. Considering this task on its own, then, there was no advantage to be gained in strategically directing attention to either location prior to the onset of the probe – the implication being that the observed attentional bias towards predictive stimuli did not reflect strategic allocation of attention, but rather an involuntary process. The long SOA condition may then have provided sufficient time for a more strategic, top-down attentional process to return attention to the centre of the display.

However, an alternative account is possible. Notably, the dot probe task was embedded within predictive learning trials in this experiment, and this overlap in task structures raises questions over the strategies that participants might have used. In particular, while participants were instructed to ignore the stimuli until after they had responded to the dot probe, they may nevertheless have begun a strategic process of identifying the stimuli and preparing a categorization response prior to the onset of the probe. On this account, then, the rapid attentional bias towards predictive stimuli demonstrated in the dot probe task may result from a voluntary process. The absence of a bias at long SOA might then be because 1000ms provided sufficient time for participants to program a categorization response and then return attention to the centre of the display in anticipation of the upcoming dot probe. Additionally the fact that RTs in the short SOA group were longer than in the long SOA group may also be seen as consistent with the idea that participants spent time preparing for a categorisation response before responding to the dot. According to this, the effect of SOA on RTs may be seen as a typical case of cognitive bottle neck in concurrent multitasking preparations (see [27], for a review on this issue). Note, however, that this effect of SOA on participants’ RTs has also been found even when the learning and the dot probe tasks take place in separate trial blocks [16].

Thus we have two alternative accounts: One which invokes opposing involuntary and strategic attentional processes, and the other in which allocation of attention is entirely strategic. The current findings do not allow us to decide between these alternatives (though we note that influences of experienced predictiveness on dot probe performance can be observed even when the two tasks are entirely separate, which is harder to reconcile with the wholly-strategic account; see Experiment 2 in [16]). For current purposes this issue is not critical, however: The important finding is that the processes underlying the influence of learned predictiveness on attention show distinct influences of selection history and explicit knowledge. This is true whether we align selection history with involuntary and explicit knowledge with voluntary attention, or whether selection history and explicit knowledge both exert distinct effects on strategic orienting. Having established a distinction here, future studies could further investigate the nature of the underlying cognitive processes.

